# Beyond the harem: vocal complexity in a captive gelada (*Theropithecus gelada*) bachelor group

**DOI:** 10.64898/2026.09.01.748493

**Authors:** Alma M. N. Nederlof, Jeroen Kappelhof, Anne Marijke Schel

## Abstract

The social complexity hypothesis for communication posits that species with elaborate social systems require more sophisticated vocal communication to navigate social life. Geladas (*Theropithecus gelada*), who live in complex multilevel societies and form diverse social bonds, are ideal for testing this hypothesis. Males produce species-specific, or ‘derived’, calls that reflect social status and function in facilitating male-female bonding. However, although derived call use also occurs in male-male social interactions, their function in male–male communication remains unexamined. In this pilot study we investigate the use of derived calls within a captive all-male bachelor group and test two hypotheses: (1) derived calls signal bachelor dominance status, and (2) derived calls function to maintain male–male social bonds. Despite a limited sample size, our results show that bachelor leaders produce higher proportions of derived calls than bachelor followers, mirroring patterns found in harem groups. Furthermore, bachelor followers were most likely to use derived calls in affiliative and resting contexts, suggesting a role in social bonding. In contrast, bachelor leaders used them more broadly. They additionally produced derived calls during agonism and travel, thereby possibly reflecting their more elaborate leadership duties. These findings provide the first preliminary evidence that derived calls serve functions beyond male–female communication and may be integral to both status signalling and affiliative interactions among bachelor males. Our results on the vocal dynamics in male geladas shed further light on the co-evolution of social and vocal complexity and offer broader implications for the origins of human language.

## 1. Introduction

Different animal species show varying levels of complexity within their vocal communication. A possible evolutionary explanation for this variety lies in the differences found in their social systems. This idea is formalised in the *social complexity hypothesis for communication* (SCHC), which proposes that species with higher levels of social complexity require more elaborate communicative behaviour; a concept also thought to underlie the evolution of human speech and language (Freeberg et al. 2012).

Social complexity results from the dynamic interplay between the core components shaping an animal species’ social system, i.e. its social organisation (e.g. group size, -composition, and -cohesion), its social structure (e.g. the patterning of social interactions), its mating system, and its care system (Kappeler 2019). For example, repeated affiliative or agonistic interactions within a group setting can lead to the formation of social bonds and specific types of dominance hierarchies (Kappeler 2019). These two social strategies require specialized socio-cognitive adaptations and become more complex to manage depending on the social dynamics and size of the group (Seyfarth & Cheney 2015; Byrne 2016). Vocal complexity, on the other hand, is typically defined by repertoire size (i.e. the number of distinct call types), but may also include how call types are organized into sequences, their acoustic variation, and contextual flexibility (Bouchet et al. 2013; Freeberg et al. 2012).

Support for the SCHC comes from several comparative studies across primates (Bouchet et al. 2013; McComb & Semple 2005), birds (Freeberg 2006; Krams et al. 2012), bats (Knörnschild et al. 2020), and mongooses (Manser et al. 2014), which consistently show a positive correlation between the previously mentioned forms of social complexity and vocal complexity. Yet, despite it being another critical part in understanding the co-evolution of social and vocal complexity, how vocal complexity actually *functions* within socially complex species still remains relatively poorly understood (Gustison et al. 2012; 2019).

A suitable species for studying this are geladas (*Theropithecus gelada*). Geladas are the closest living sister lineage to the baboons of the *Papio* genus within the *Papionini* tribe of cercopithecine monkeys. They form a distinct monotypic genus that diverged from the common ancestor about 4.7 mya, but still shares clear morphological, ecological, and behavioural traits with some of the closely related baboon species (Hammerschmidt & Fischer 2019; Maciej et al. 2013; Zinner et al. 2018). Geladas are endemic to the Ethiopian highlands and the only graminivorous primate, making their ecological complexity fairly low (Fashing et al. 2014). Concerning their social complexity, geladas form multi-level societies in the wild in which smaller social units are embedded within progressively larger social constructs.One of the smallest units isthe reproductive unit (mean size of 12.2; Dunbar 1980) comprised of a dominant ‘leader’ male, 1-11 adult females with their offspring, and occasionally 0-3 subordinate ‘follower’ males (Kawai et al. 1983; Snyder-Mackler et al. 2012b). Multiple reproductive units (5-30) form bands that forage and sleep together (Kawai et al. 1983). Associated with these bands are all-male bachelor groups (mean size of 4.7; Pappano 2013) that are composed of adolescent males that have dispersed from their natal unit (Kawai et al. 1983) and former leader males (Pappano 2013). Lastly, an additional higher form of organisation in gelada society are the communities, that may consist of more than 100 units that are loosely associated (Kawai et al. 1983; Snyder-Mackler et al. 2012b).

Within this multilevel society, both cooperation and competition for male leadership in reproductive units shape male-, and partly female-, behaviour. In particular, leader males living *within* reproductive units often form long-term affiliative bonds with their females (Gustison et al. 2012), a strategy likely enhancing their reproductive success. In contrast, males living *outside* of reproductive units typically spend three to five years in bachelor groups before potentially gaining reproductive opportunities. This can be achieved by challenging a harem leader and taking over his reproductive unit (Dunbar 1984; Pappano, 2013). These takeovers can occur cooperatively and are usually preceded by ritualised vocal contests where bachelors and reproductive unit leader males exchange loud display calls (Benítez et al. 2016), comparable to those found in the closely related chacma baboons (*Papio ursinus*; Fischer et al. 2004). These display calls signal male quality (Benítez et al. 2016), which bachelors use to target lower-quality leaders for takeovers (Benítez et al. 2017). At the same time, female behaviour can also influence takeover outcomes. Unlike the behavioural patterns found in other closely related species, e.g. the hamadryas baboons (*Papio hamadryas*), where leaders aggressively control female movement (Swedell & Schreier 2009), female geladas may collectively either support the leader male or abandon him in favour of the challenger (Dunbar 1984). This re-emphasises the importance for leader males to maintain good relationships with his females. Thus, with takeovers playing a central role, geladas show diverse social dynamics that define their social relationships and, by extension, their social complexity (Kappeler 2019).

Consistent with the SCHC, geladas show vocal complexity alongside their social complexity. In addition to the basic vocal repertoire of call types shared with all *Papio* spp. (Hammerschmidt & Fischer 2019), the gelada repertoire is expanded through the inclusion of unique *derived* (i.e. species-specific) calls. These include inhaled grunts, exhaled and inhaled moans, exhaled and inhaled wobbles and vocal yawns (Table 1; Figure 1), which are often recombined in sequences with (non-derived) exhaled grunts (Gustison et al. 2012). These sequences are primarily produced by adult males, who commonly use them in short-range social interactions with females (Gustison et al. 2012; Gustison & Bergman 2016). Due to their acoustic characteristics, such as frequency modulation and rapid alternation of inhaled and exhaled forms, producing such sequences including derived calls requires high muscular effort of the lungs to produce them loud enough. Hence, sequences including derived calls are generally considered energetically costly and complex to produce (Bergman 2013; Gustison et al. 2016; Gustison & Bergman 2017; Titze & Riede 2010). The presence of such complex calls and sequences in the gelada vocal repertoire suggest an important adaptive function (Bergman 2013; Gustison et al. 2016). Notably, the use of similar types of derived vocalisations are absent in those *Papio* spp. that, like geladas, live in multilevel societies (i.e. Guinea baboons (Papio papio) and Hamadryas baboons; Gustison et al. 2012; Hammerschmidt & Fischer 2019; Maciej et al. 2013). Taken together, this raises questions about the potentially unique social pressures that have selected for the evolution of these calls in geladas, as well as their exact function in gelada society.

**Figure 1.**
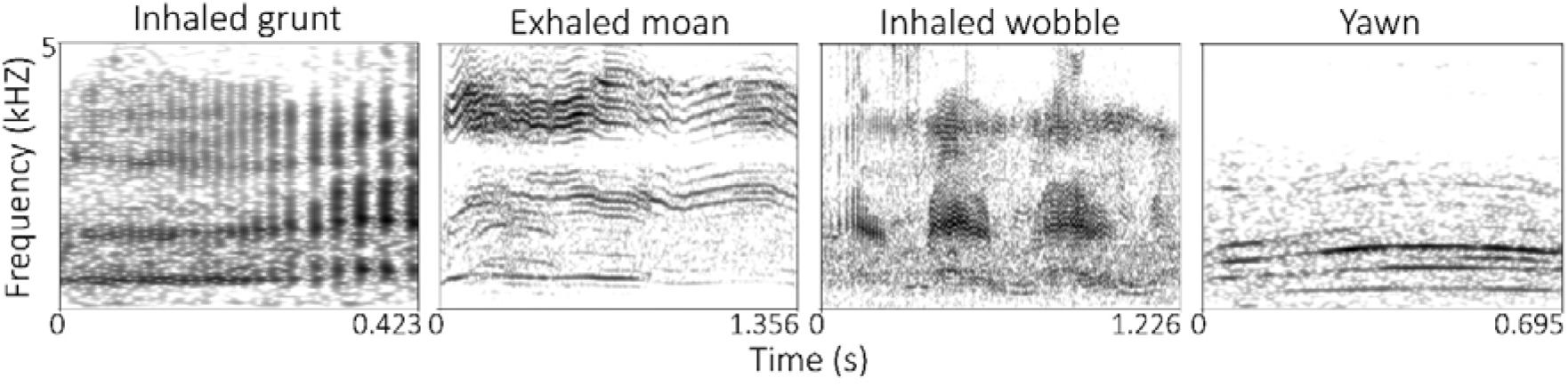
Spectograms of the derived call types produced by the gelada’s in this study (made in Praat: Boersma & Weenink 2024).

**Table 1.**
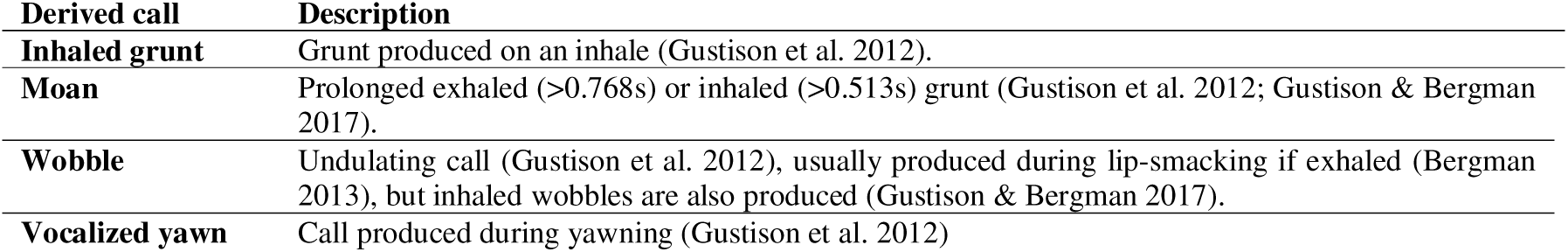
Description of the derived call types gelada males combine in a vocal sequence with non-derived grunts during social interactions

So far, the use of derived calls and derived call sequences has mainly been studied within reproductive units, where they fulfil several functions. First, the frequency of complex sequences (i.e. those including at least one derived wobble, moan, or yawn vocalization) increases with conspecific noise, likely aiding within-unit communication when joining in larger groups, as variation in vocalisations could help overcome noise (Hotchkin & Parks 2013; Gustison et al. 2019). Second, playback experiments revealed that females are more attentive to complex over simple sequences (i.e. without derived calls), suggesting a possible role in sexual selection and male quality signalling (Gustison & Bergman 2016). This is reinforced by the finding that males of higher status (i.e. harem leader males), who are likely of higher quality (Benítez et al., 2016), produce more derived calls than males of lower quality (i.e. follower males; Gustison et al. 2019). Third, derived calls facilitate male–female bonding, with complex sequences often prompting females to groom their leader male (Gustison et al. 2019). And finally, captive studies have shown that reproductive unit leader males emit derived calls towards their females after conflicts within their unit, possibly aiding reconciliation and consolation that promote bond repair and restore social cohesion within the unit (Gustison et al. 2012; Leone & Palagi 2010; Palagi et al. 2018; but see Hohn et al. 2024 where no post-conflict repair behaviour was found in wild geladas). However, while the aforementioned studies have mainly examined the role of derived calls in male-female social interactions, it currently remains unclear whether they function exclusively in male-female social interactions or also play a role in male-male social interactions within bachelor groups (Gustison et al. 2019).

This is surprising, as previous research has shown that social relationships between gelada bachelors are more complex than previously assumed (Pappano 2013), and the use of derived calls in male-male interactions is not uncommon (pers. obs.). In this study we therefore investigated the possible functions of derived calls within bachelor male gelada groups.

There are two possible non-mutually exclusive functions for the use of derived calls in bachelor groups, which are both related to the proposed functions of derived calls in reproductive units: signalling of male status (and quality), and the formation and maintenance of male-male bonds. First, derived calls may function in the assessment of male quality and status and thereby contribute to the formation of dominance hierarchies in bachelor groups. This is relevant in the context of male takeovers, as the male gaining the leader position after a cooperative takeover will ultimately have the highest reproductive success (Snyder-Mackler et al. 2012a). However, which bachelor male becomes the reproductive unit leader after a cooperative takeover is poorly understood. Dominant males, possessing specific characteristics indicating male quality and fighting ability likely have a greater chance. Besides morphological traits enhancing direct fighting capabilities, such as larger canines (Galbany et al. 2015) and greater body size (Wright et al. 2019; Plavcan 2012), these may also include more indirect cues, such as vocalizations containing information about the quality and competitive capability of an individual (e.g. Benítez et al. 2016; Fitch 1997; Fitch & Hauser 2003; Puts et al. 2016). Indeed, in geladas, the earlier described non-derived loud display calls, produced between reproductive unit leader males and bachelor males prior to takeover attempts, contain acoustic parameters linked to male quality and status (Benítez et al. 2016). Yet, whether derived calls, previously implicated as potential indicators of male quality in the context of female choice (Gustison & Bergman 2016), also play a role in the assessment of male quality during short range interactions within bachelor groups, and in the potential formation of dominance hierarchies in these groups, is currently unknown.

Secondly, derived calls could contribute to forming and maintaining bonds. Alongside the inevitable competition between bachelor males for a future harem leader position, males within bachelor groups benefit from forming social bonds with other bachelors, as such bonds help secure coalitionary support during cooperative takeovers of reproductive units (Dal Pesco et al. 2022; Pappano 2013). Although not all coalition partners can become leader males after a takeover, males that participated in a takeover alongside the new harem leader male are usually allowed to join the reproductive unit as follower males (Dunbar 1984; Snyder-Mackler et al. 2012a). This is beneficial for both parties, as leader males with follower males have approximately 30% longer tenures due to increased protection from other bachelors, and follower males may get to sire some offspring (Snyder-Mackler et al. 2012a). Accordingly, previous research has shown that bachelor males spend significant time grooming each other, more than adult females do (Pappano 2013). However, no study has yet investigated whether bachelor males also use derived calls to strengthen their social bonds, as seen in male–female bonding (Gustison et al. 2019).

Building onto the previous work on derived call use and its form and function in male-female social interactions (Gustison et al. 2012; 2019), in this study we hypothesized that, similar to how they function in male-female interactions, derived calls in bachelor groups serve a dual function, i.e. i) indicating male quality and asserting dominance status and ii) forming and maintaining male-male bonds. If i) derived calls function to indicate male quality and assert dominance status, we expected higher-ranking males to produce a higher frequency of complex vocal sequences as well as a higher proportion of derived calls per (complex) sequence compared to lower-ranking males. If ii) derived calls function to maintain male-male bonds, we expected the likelihood of producing complex sequences to be higher in affiliative contexts compared to other contexts.

In addition, we aimed to broadly compare the vocal behaviour (i.e. frequency of derived calls and production contexts) between bachelor males and reproductive unit leader males to explore potential parallels in the use of derived calls. Such parallels could be informative because both harem leaders and dominant bachelor rely on maintaining bonds within their respective groups: harem leaders must maintain bonds with females to avoid abandonment during takeover attempts (Dunbar 1984), while dominant bachelor males may require bonds with other bachelors for coalitionary support to achieve successful takeovers (Pappano 2013). If both types of males use derived calls to maintain bonds, this would suggest that male-male social relationships are more intricate than previously recognised, adding another, formerly unregistered, level of complexity to this species’ social system.

## 2. Methods

### 2.1. Subjects & location

Observations were made in November and December 2023 at Rotterdam Zoo, The Netherlands. Seven gelada males were observed (Table 2). Six of these males were housed together in a bachelor group in an outdoor enclosure of 755 m^2^, adjacent to the outdoor enclosure of the reproductive unit (865 m^2^; S. Lughthart, pers. comm., 2025; see Supplementary Material 1, Figure S1 & S2 for maps of the enclosures). The reproductive unit consisted of one leader male, four females and their offspring. Both outdoor enclosures contain shrubs and climbable rock structures. The substrate for the reproductive unit was mainly sand, whereas that of the bachelors was primarily soil and also included a small water stream. A central building between both outside enclosures contained the visually separated indoor enclosures of both groups, with an outdoor visual barrier on its roof preventing the groups from seeing, but not hearing, each other. The indoor enclosure of the reproductive unit (90 m^2^) featured rock-textured walls and flooring. The indoor bachelor enclosure, with straw bedding, and the caretakers’ area (73 m^2^; S. Lugthart, pers. comm., 2025) were not accessible to the public or researchers. The animals had access to the indoor areas at all times, except during keepers’ cleaning hours. They were fed vegetables and hay daily, with occasional provision of monkey pellets and fresh grass. No fixed herbivorous substrate was present in the enclosures that allowed for unprovisioned foraging.

**Table 2.** The names of the study subjects, the group they belonged in and the age class of the individuals (Dunbar 1980)

| Name (abbreviated) | Group | Age class |
| --- | --- | --- |
| **JB** | Reproductive unit | Adult |
| **HG** | Bachelor | Adult |
| **ST** | Bachelor | Adult |
| **DP** | Bachelor | Young adult |
| **LP** | Bachelor | Young adult |
| **DE** | Bachelor | Subadult |
| **DO** | Bachelor | Subadult |

One important note to make here is that during data collection it already became clear that the bachelor group had seemingly divided itself into two subgroups, as certain bachelors were rarely observed interacting with each other. Indeed, this observation was later confirmed when logging the vocalisations and their contexts into excel (see below). For example, of all single vocalisations and sequences recorded during social interactions in the bachelor group, only 20/1081 (1.9%) occurred between individuals of the two subgroups. Consequently, we further considered the bachelor group to consist of a subgroup with the individuals HG, LP and DE, and a second subgroup consisting out of ST, DP and DO. Small bachelor groups are also a regular occurrence in the wild, with an average size of 4.7 individuals (Pappano 2013).

### 2.2. Data collection

Each individual was observed during independent 15m focal observation bouts for three to four times per week (Altmann 1974), until approximately 5 hours per individual were recorded (total = 2084.5min; range = 282.5 – 306 min per individual). Thus, blind recording of the data was not possible, because the study involved identifying individuals during focal observations. Observation sessions were conducted between 9:30 am and 3:30 pm, following a randomized daily order. During focal observations, continuous video and audio recordings were made simultaneously. Videos were recorded with a JVC Everio R camcorder and vocalisations were recorded with a Marantz PMD660 solid state recorder connected to a Sennheiser MKE600 directional microphone. Individuals were observed at the edge of the enclosures where the visibility was highest. This was at a maximum distance of 25 meters, which allowed the directional microphone to adequately pick up vocalisations. Whenever any individual was observed vocalising during these observations, the individual who vocalized, and the production context were spoken into the recording. In this way, if needed, also vocalisations from non-focal individuals could be used in further vocal sequence analyses (see below). Focal observations were stopped when individuals went inside, and later finished.. In addition, ad libitum audio recordings (total = 674min) were performed in situations where individuals that were already focally observed were the only ones in sight. These ad libitum observations increased the final dataset used for analyses at the sequence level. These were stopped again if a planned focal individual would come back in sight.

### 2.3. Data analysis

#### 2.3.1. Determining dominance ranks

To assess the association between derived call production and dominance rank, we first determined the dominance rank of the bachelor males. All focal video recordings were coded with ZooMonitor (Lincoln Park Zoo 2022) to extract frequencies of dominance (i.e. displacement) and submissive (i.e. giving ground, bare teeth; Supplementary Material 1, Table S1) behaviours of focal individuals. Elo scores were then calculated in R (version 4.4.0, R Core Team 2022) using aniDom (Farine & Sanchez-Tojar 2021), with initial scores set at 1000 and a K-value of 100, following Tinsley Johnson et al. (2014). The K-value is the number of points a male gains after winning (i.e. dominance) or losing (i.e. submission) the interaction, and additionally, these interactions are weighted by predicted win-loss probabilities (Neuman et al. 2011; Tinsley Johnson et al. 2014). Behavioural coding from focal video recordings, and vocalization contexts (see below for details) from both the non-focal and ad libitum audio recordings provided the data for the dominant and submissive behaviours.

Because, as mentioned, preliminary analyses revealed that the behavioural and vocal interactions between the subgroups were rare, and without any clear winners or losers, the dominance ranks of the bachelors were calculated within their respective subgroups. This was considered a valid solution, as Elo scores remain suitable for calculating the dominance rank in groups with a small number of individuals (Neumann et al. 2011). Nonetheless, to evaluate the robustness of these hierarchies, we estimated uncertainty using the splitting method. Within the two subgroups, the ordering of interactions was randomised 1000 times and from this, correlations were calculated between Elo scores derived from halved datasets, with a correlation above 0.5 suggesting a robust dominance hierarchy (Farine & Sanchez-Tojar 2021).

#### 2.3.2. Vocal analysis

All audio recordings collected during focal and ad libitum observations were saved as WAV files. Recordings were played back in Audacity (The Audacity Team 2024) to identify single vocalisations and vocal sequences. For each sequence, the caller(s), behavioural context, and any spoken notes, were extracted and logged in Excel. Sequences in which the caller could not be reliably identified were discarded. For each focal individual, we then identified the number of vocal sequences produced during their total focal observation time (5 hours per individual). A sequence consisted of two or more non-derived and/or derived calls, separated from other sequences by a silence of at least 2.98s (Gustison et al. 2019).

Sequences were assigned to one of five context categories: affiliation, agonistic, foraging, rest or travelling (Supplementary Material 1, Table S1). These categories follow Gustison et al. (2019), however, agonistic was added as a context to account for the potential function of derived calls in competitive contexts (Gustison et al. 2012; Leone & Palagi 2010). Vocalisations produced by the reproductive unit leader male during copulation (50s of total focal time) were excluded for fair comparison with bachelor males. Sequences were also excluded when the context category could not be reliably determined, when caretakers were involved, during display call interactions, or when calls were inaudible due to high background noise.

To subsequently determine whether and how the males used complex sequences (i.e. containing at least one derived call; Gustison et al., 2019) depending on rank and context, we determined the amount of complex versus simple sequences, as well as the proportion of derived calls per sequence. For this, we simply counted the number of derived and non-derived vocalisations per sequence. Because of the clear acoustic differences across derived calls and non-derived calls, the classification of call types was mostly done by ear and visual inspection of the spectrogram. However, if call types were unclear, the spectrogram was inspected for further acoustic patterning and details in Praat (Boersma & Weenink 2024, see also Figure 1). Because moans are defined as elongated grunts, they were distinguished based on call duration thresholds (cf. Gustison & Bergman 2017; Table 1). The counts of derived and non-derived calls were used to classify a sequence as complex (i.e. containing at least one inhaled grunt, moan, wobble or yawn) or simple (i.e. containing only exhaled grunts), and to compute proportions per sequence for further statistical analyses. Contrary to Gustison et al. (2019), we included inhaled grunts as part of complex sequences, due to their complex nature (i.e. low in frequency and part of an exhale-inhale vocal pattern; Titze & Riede 2010; Gustison & Bergman 2017; Gustison et al. 2012; Hewitt et al. 2002; pers. obs.).

### 2.4. Statistics

Generalized linear mixed models (glmm’s) were used to investigate the relation between bachelor male vocal sequence complexity, dominance rank, and production context. First, we investigated whether the likelihood of producing a complex sequence instead of a simple sequence differed depending on rank. Because this analysis concerns the rate at which individuals produced different sequence types, only sequences recorded during focal observations were included. Any potential bias would arise by including ad lib data, for example, individuals who vocalised more or were more often present outside, were prevented by this measure. In this first glmm, with a binomial family, the type of sequence was taken as the response variable (binary; 0 = simple, 1 = complex), rank as fixed factor, and vocalising individual as random factor. For the next analyses, all recorded sequences (i.e. recorded during both focal and ad libitum observations) were used, These analyses focused on properties of the sequences themselves (e.g. the proportion of derived calls within a sequence and the contextual use of sequences), rather than on individual production rates. Including all available sequences increased the sample size substantially and therefore the reliability of the analyses. In the second model, in order to compare results with Gustison et al. (2019), we next tested differences in the proportion of derived calls per sequence produced by the bachelor males of different rank. In this second model, with a beta-binomial family, the response variable was a cbind with the number of derived calls and non-derived calls per sequences, the fixed factor was rank, and the random factor was the vocalising individual. This same model was used with a subset of data with only complex sequences, to test whether potential statistical differences found in the previous model would not just be an artifact of different ranks producing different amounts of complex sequences. This analysis could additionally identify whether not only the frequency of complex sequence production but also the sequence composition differed depending on status. Lastly, to gain insight into the likelihood of producing a complex sequence in the different social contexts, and whether this likelihood differed depending on rank, a final model with a binomial family was used, with the type of sequence as the binary response variable (i.e. 1 = complex sequence, 0 = simple sequence), the interaction status*context as fixed factor, and vocalising individual as random factor.

The glmmTMB package (Brooks et al. 2017) in R (R Core Team 2024) was used for performing these models. ggeffects (Lüdecke 2018) was used to extract predicted values from the glmm’s. To test if the models met the assumptions, the DHARMa package (Hartig 2022) and the performance package (Lüdecke et al. 2021) were used. The post-hoc pairwise comparisons of the last model, that investigates the likelihood of a complex sequence per context and per status, were adjusted using Tukey’s method (Agbangba et al. 2024) via the emmeans package (Lenth 2024).

The reproductive unit leader male was excluded from the statistical analyses due to him being only one male. Instead, descriptive statistics were used to compare the behaviour of bachelor males to that of the reproductive unit leader male. Statistical and descriptive plots were constructed using the ggplot2 package (Wickham 2016).

## 3. Results

### 3.1. Relationship between dominance and derived call production

To determine if and how derived call production was related to dominance, we first calculated the Elo scores from the coded dominance interactions. A total of 88 dominance interactions (including displacement, give ground and/or bare teeth) were recorded during focal observations (2.5 dominance interactions per hour). More severe forms of aggression, such as chases and bidirectional fights, were rare within the subgroups and were both only recorded once. The Elo scores for the bachelor individuals showed one consistently high-ranking male in both subgroups, followed by two males with similarly lower scores (Table 3). Dominance interactions among the lower-ranking males were rare, and consequently, most dominance information came from encounters between the top-ranking male and the subordinates. This pattern indicates a clear leader in each subgroup, with a weak hierarchy among the remaining individuals. Accordingly, each bachelor male was therefore classified as either “leader” or “follower”, consistent with classifications used for males in reproductive units (Gustison et al. 2019). Bachelor leaders included both an adult and a young adult, while followers spanned all age classes (Table 3). Estimating uncertainty by splitting (Farine & Sanchez-Tojar 2021) yielded mean correlations of 0.767 for subgroup I and 0.680 for subgroup II, indicating that the dominance structure was robust despite the small subgroups.

**Table 3.** The Elo scores and subsequent status (BL = bachelor leader, BF = bachelor follower, and RUL = reproductive unit leader) given to the different subjects of different age classes (Dunbar 1980), along with an overview of the total calls (both single vocalisations and vocalisations contained in sequences) and total number of sequences individuals produced during their respective five hour focal observations (not including ad libitum recorded vocalisations).

| Individual | Age class | Elo score | Status | #Derived calls<br>(total #calls) | #Complex sequences<br>(total #sequences) |
| --- | --- | --- | --- | --- | --- |
| *Subgroup I* |  |  |  |  |  |
| **HG** | Adult | 1266.34 | BL | 150 (408) | 50 (65) |
| **LP** | Young adult | 909.46 | BF | 18 (139) | 11 (41) |
| **DE** | Subadult | 824.21 | BF | 22 (157) | 9 (34) |
| *Subgroup II* |  |  |  |  |  |
| **DP** | Young adult | 1290.01 | BL | 89 (370) | 32 (71) |
| **ST** | Adult | 889.75 | BF | 10 (109) | 7 (32) |
| **DO** | Subadult | 820.24 | BF | 14 (160) | 10 (35) |
| **JB** | Adult | - | RUL | 62 (237) | 24 (41) |

In general, bachelor leaders and the reproductive unit leader produced more calls than bachelor followers (Table 3). Correspondingly, bachelor leaders had the highest derived call rates (HG: 30.1 derived vocalisations per focal observation hour; DP: 18.9 derived vocalisations per hour), followed by the reproductive unit leader (JB: 12.7 derived vocalisations per hour). The bachelor followers had the lowest derived call rates (LP: 3.6 derived vocalisation per hour; DE: 4.5 derived vocalisations per hour; ST: 2.0 derived vocalisations per hour; DO: 2.8 derived vocalisations per hour; Figure 2).

**Figure 2.**
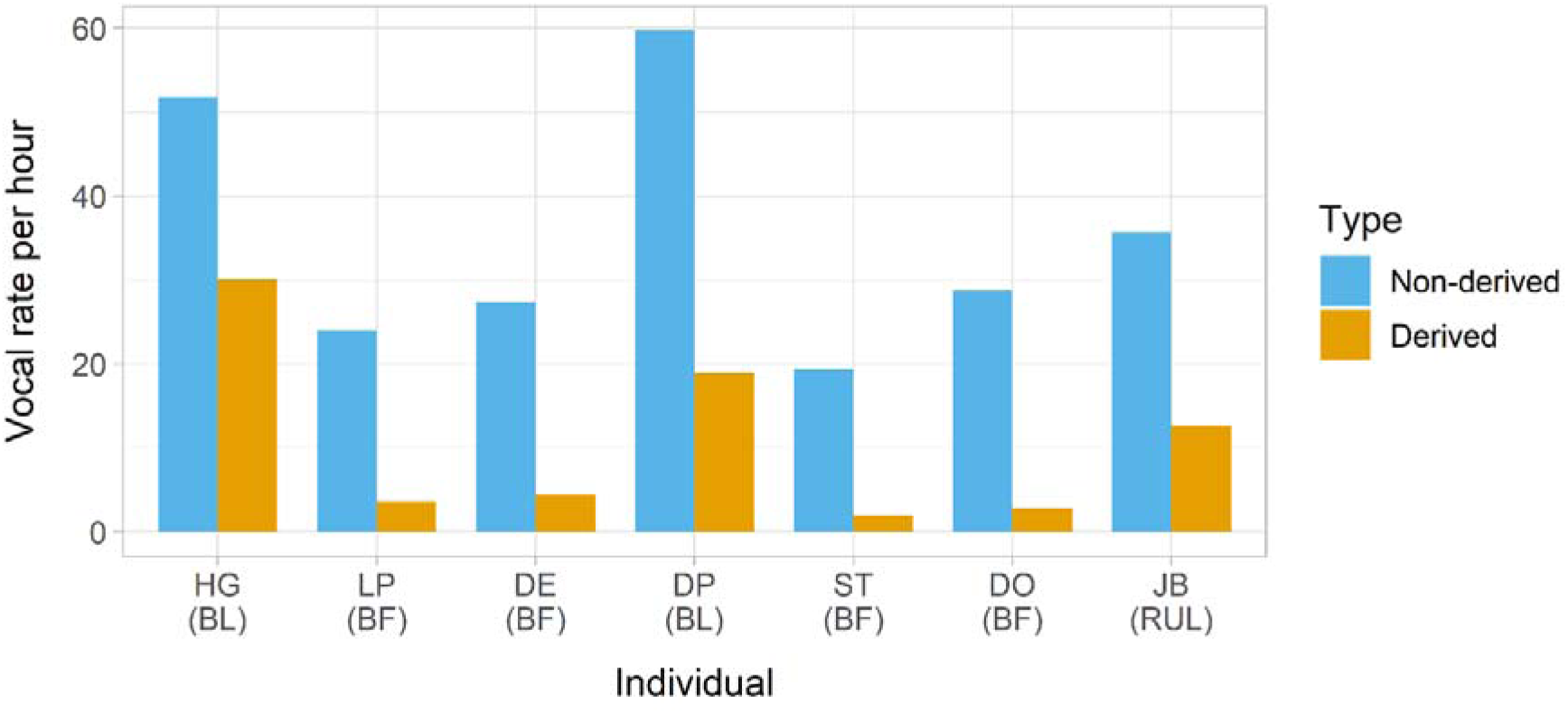
The non-derived and derived call rate per hour for each individual, calculated from vocalisations individuals produced during their respective focal observations only (see Table 3). BL = bachelor leader; BF = bachelor follower; RUL = reproductive unit leader

Bachelor leaders were significantly more likely to produce complex sequences (predicted probability = 0.61, 95% CI = 0.46 – 0.75) than bachelor followers (predicted probability = 0.26, 95% CI = 0.17 – 0.37; *z* = 3.641, *p* = 0.0003, *n* = 278 sequences produced by focal individuals during their respective focal observations). In addition, the bachelor leaders and reproductive unit leader produced higher proportions of complex sequences compared to the bachelor followers Table 3; Figure 3).

**Figure 3.**
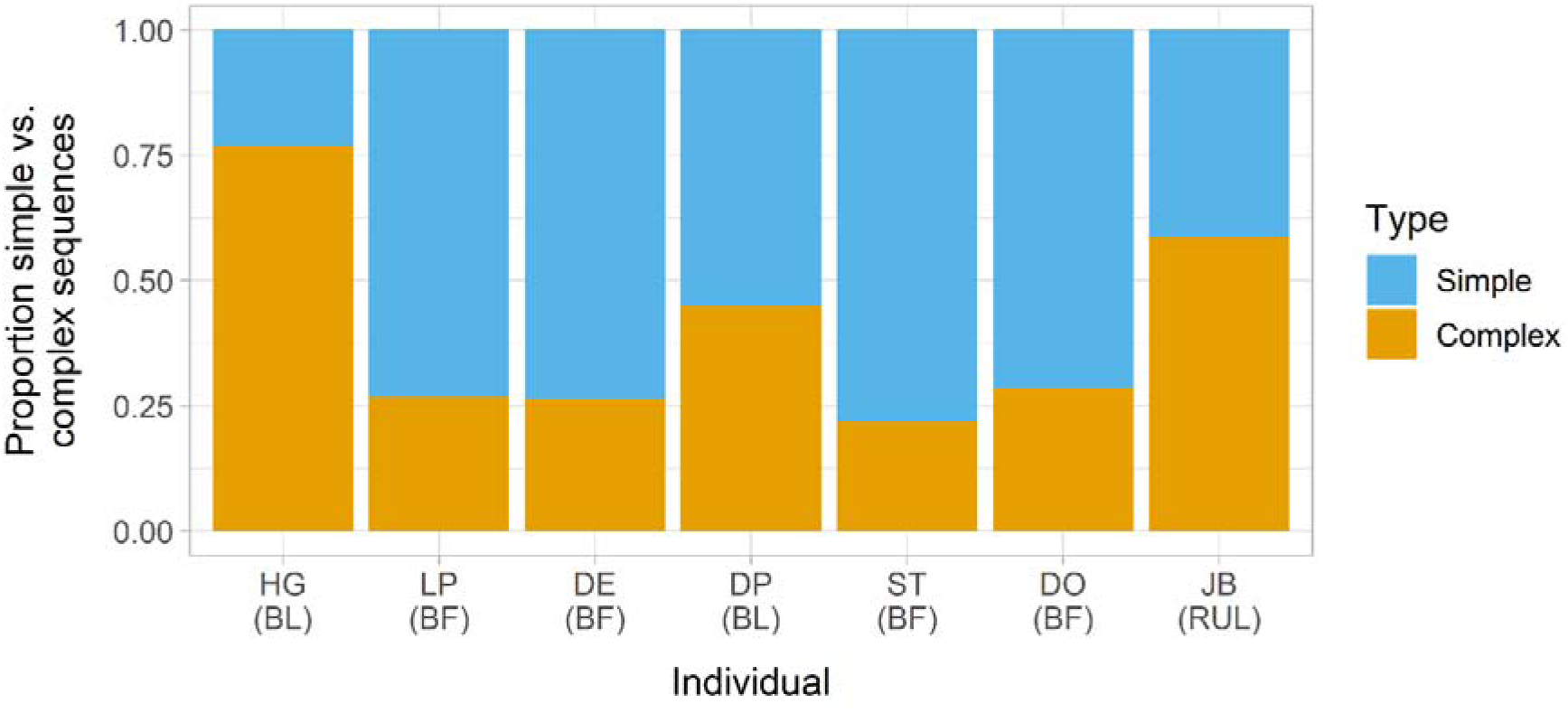
The proportion of simple sequences (i.e. containing no derived calls) and complex sequences (i.e. containing one or multiple derived calls) each individual produced during their five hour focal observations only. BL = bachelor leader; BF = bachelor follower; RUL = reproductive unit leader

When further looking into the overall composition of all sequences collected during both focal and ad libitum observations, followers in bachelor groups used a significantly lower proportion of derived calls within their sequences (predicted proportion of derived calls per sequence = 0.11, 95% CI = 0.08 – 0.14) compared to the leaders in bachelor groups (predicted proportion of derived calls per sequence = 0.22, 95% CI = 0.16 – 0.29; *z* = -3.200, *p* = 0.0014, *n* = 1053 sequences, Figure 4a). In further descriptive analyses that also included the reproductive unit leader, both the reproductive unit leader (*M ± SE* = 0.167 ± 0.017, *n* = 125 sequences) and bachelor leaders (*M ± SE* = 0.178 ± 0.009, *n* = 562 sequences) produced higher proportions of derived call types per sequence than bachelor followers (*M ± SE* = 0.091 ± 0.008, *n* = 491 sequences; Figure 4b).

**Figure 4.**
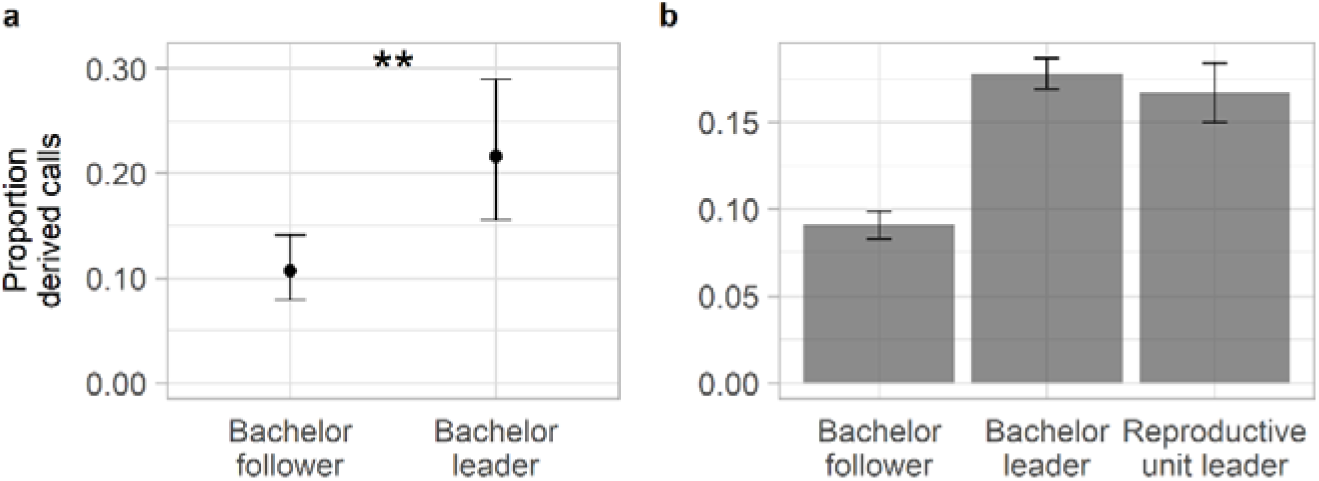
Comparisons of the proportion of derived calls contained in sequences, per status. a) predictive output of the glmm testing differences in the proportion of derived calls per sequence produced by followers and leader within the bachelor group. Points and whiskers represent the predicted mean and 95% CI. ** = *p* < 0.001. b) the observed proportions of derived calls per sequence produced by bachelor leaders, bachelor followers and the reproductive unit leader. Bars and whiskers represent M ± SE

Finally, we tested whether bachelor leaders and followers differed in derived call proportion when only taking into account complex sequences in these analyses. Here, bachelor leaders (predicted proportion = 0.32, 95% CI = 0.24 – 0.33) and bachelor followers (predicted proportion = 0.28, 95% CI = 0.27 – 0.38) did not differ statistically (*z* = 1.016, *p* = 0.3096, *n* = 439 complex sequences). Descriptively, the reproductive unit leader had a similar proportion of derived calls in complex sequences (M ± SE = 0.29 *±* 0.02, *n* = 73 complex sequences) compared to bachelor leaders (M ± SE = 0.33 *±* 0.01, *n* = 299 complex sequences) and bachelor followers (M ± SE = 0.32 *±* 0.01, *n* = 140 complex sequences).

### 3.2. Contextual usage of derived calls

When looking into the contextual usage of derived calls, we found that, overall, most vocal sequences were produced during affiliative contexts (*n* = 391/1178 sequences (33.2%); 213 simple and 178 complex), followed by the foraging (*n* = 300/1178 sequences (25.5%); 189 simple and 111 complex), travelling (*n* = 267/1178 sequences (22.7%); 164 simple and 103 complex), rest (*n* = 144/1178 sequences (12.2%): 65 simple and 79 complex), and agonistic contexts (*n* = 76/1178 (6.5%); 35 simple and 41 complex).

When subsequently assessing the distribution of complex sequence use per call context *across* the different bachelor status categories, we found that bachelor followers were overall significantly less likely to produce complex sequences than bachelor leaders (affiliation: β ± SE = 0.477 *±* 0.168, *z* = -2.098, *p* = 0.0359; agonistic: β ± SE = 0.132 *±* 0.102, *z* = -2.622, *p* = 0.0087; foraging: β ± SE = 0.279 *±* 0.109, *z* = -3.261, *p* = 0.0011; rest: β ± SE = 0.327 *±* 0.152, *z* = -2.399, *p* = 0.0164; travelling: β *± SE* = 0.171 *±* 0.069, *z* = -4.354, *p* < 0.0001; Figure 5a).

**Figure 5.**
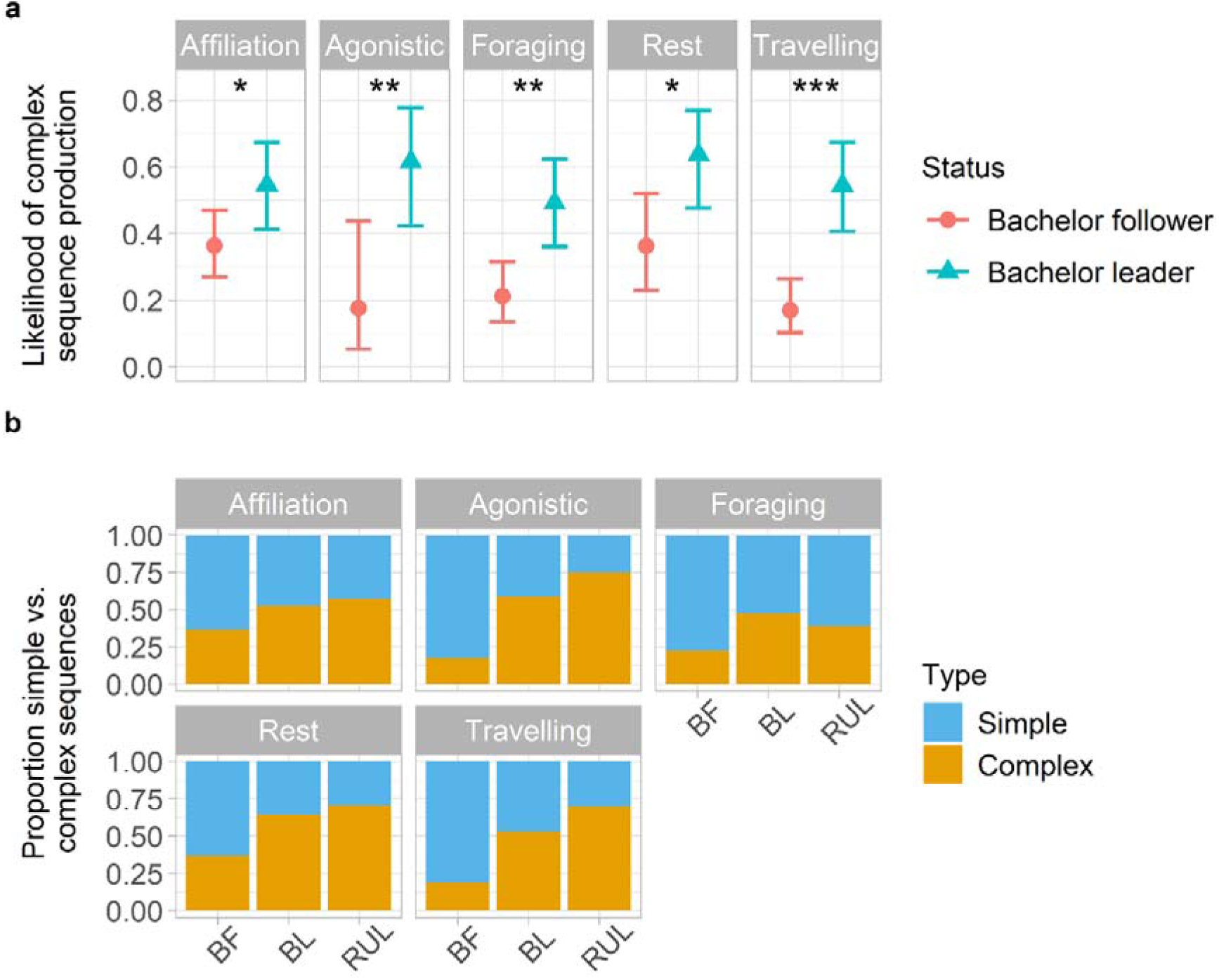
Comparisons of the likelihood of producing a complex sequence per context and per status. **a)** predictive output of the glmm model testing differences in the likelihood of producing a complex sequence per context and per status, within the bachelor group. Points and whiskers represent the predicted mean and 95% CI. * = *p* < 0.05, ** = *p* < 0.001, *** = *p* < 0.0001. **b)** observed proportions of complex sequences vs. simple sequences produced per context and status. BF = bachelor follower; BL = bachelor leader; RUL = reproductive unit leader

In corresponding descriptive analyses, the reproductive unit leader produced the highest proportion of complex sequences in the agonistic contexts (75.0%, *n* = 15 complex sequences out of the total of 20 sequences the reproductive unit leader produced in that context), followed by the resting (70.6%, *n* = 12/17 sequences) and travelling (70.0%, *n* = 7/10 sequences) contexts. In the affiliative context, complex sequences still constituted over half of his sequences (57.5%, *n* = 27/47 sequences). The lowest proportion was found in the foraging context (38.7%, *n* = 12/31 sequences; Figure 5b). Bachelor leaders produced high proportions of complex sequences during the rest (64.4%, *n* = 47/73 sequences) and agonistic context (59.0%, *n* = 23/39 sequences). Similar proportions of complex sequences were produced during the travelling (53.2%, *n* = 74/139 sequences) and affiliative context (52.6%, *n* = 82/156 sequences). The lowest proportion of complex sequences was found in the foraging context (47.1%, *n* = 73/155 sequences; Figure 5b). Bachelor followers, on the other hand, only produced relatively higher proportions of complex sequences during rest contexts (37.0%; *n* = 20/54 sequences) and affiliative contexts (36.7%, *n* = 69/188 sequences), and lower proportions of complex sequences in the foraging (22.8%, *n* = 26/114 sequences), travelling (18.6%, *n* = 22/118 sequences) and agonistic contexts (17.6%, *n* = 3/17 sequences; Figure 5b).

Finally, when specifically comparing complex sequence use per call context *within* the bachelor status categories, bachelor followers were more likely to produce complex sequences during the affiliative compared to the foraging (β *± SE* = 2.129 *±* 0.588, *z* = 2.733, *p* = 0.0493) and travelling contexts (β *± SE* = 2.799 *±* 0.818, *z* = 3.522, *p* = 0.0039), as well as during the resting compared to the travelling context (β *± SE* = 2.785 *±* 1.040, *z* = 2.743, *p* = 0.0480). No further differences across any of the other contexts were found for the follower males. In contrast, bachelor leaders showed no significant differences in producing complex sequences when comparing any of the behavioural contexts.

## 4. Discussion

In this study, we investigated whether derived gelada calls, previously shown to serve multiple functions in male female communication (Gustison & Bergman 2016; Gustison et al. 2012, 2019), also function in male-male communication within bachelor groups. We hypothesized that derived calls help maintain dominance hierarchies by signalling male status and quality, and that they support male-male bond maintenance relevant for coalitionary support. Accordingly, we predicted that higher-ranking males would use derived calls more frequently than lower-ranking males, and that these calls would be more common in affiliative contexts. Although the results of our study come from two captive study groups over a two month period leading to a relatively small sample size, they largely support these predictions. Dominant bachelor leaders produced complex sequences more often than subordinate follower males and, correspondingly, a higher proportion of derived calls per sequence. This usage pattern of bachelor leaders was comparable to that of the reproductive unit leader. In addition, we found that the bachelor followers were most likely to produce complex sequences in affiliative and rest contexts. This pattern was not found in both types of leader males, who also produced complex sequences at relatively high rates in all other contexts. Despite the limited sample size, these patterns were strong and occurred across all age categories, suggesting a functional variation of derived call use depending on the males’ social roles only.

Our finding that bachelor leaders produce derived calls more frequently than bachelor follower males aligns with previous work showing that reproductive unit leaders also produce more derived calls than follower males in reproductive units (Gustison et al. 2019). One proposed function of derived calls is that females use these calls to assess male quality, thereby playing a role in sexual selection (Gustison & Bergman 2016). Indeed, complex sequence production is energetically costly, as it requires the coordination of exhaled and inhaled low- and high-frequency calls, demanding high lung pressure and precise muscular control (Gustison et al. 2016; Gustison & Bergman 2017; Hewitt et al. 2002; Titze & Riede 2010). Leader males may be able to produce more derived calls, because traits linked to dominance and quality, such as larger lung capacity and better body condition can support producing these acoustically elaborate calls at higher rates (Fitch & Hauser 1995, 2003; Puts et al. 2016; Titze & Riede 2010). Although the proportion of derived calls within complex sequences did not differ between bachelor leaders and followers, leader males did produce derived calls and derived call sequences at higher hourly rates. The frequency of derived call production may therefore function as an honest indicator of male quality beyond the context of female choice (Gustison & Bergman 2016), i.e. by facilitating male-male coalition formation and the maintenance of male dominance hierarchies without the need for physical aggression (Kavanagh et al. 2021; Tibbetts et al. 2022). However, future research should directly test links between derived call production and male quality measures such as body size, canine length, testosterone levels, and reproductive success (Fitch & Hauser 2003; Galbany et al. 2015; Plavcan 2012; Puts et al. 2016; Soma & Garamszegi 2011). In particular, studies could examine whether derived call rates predict female defection during takeover challenges and leadership outcomes following coalitionary takeovers. Moreover, besides their evolutionary significance, insights into the role of derived calls in status signalling, bachelor coalition formation, and male quality assessment may also have applied value. Specifically, zoos participating in gelada breeding programmes could use derived call production as a non-invasive behavioural indicator to determine social stability of bachelor groups and the selection of suitable harem leader males. Such applications may become increasingly important as wild gelada populations continue to decline (Gippoliti et al. 2025).

Another, not necessarily mutually exclusive, explanation for leaders producing relatively high frequencies of derived calls is that their leadership role requires vocalizing across a wider range of contexts. Indeed, bachelor leaders produced complex sequences at relatively equal rates in all contexts, whereas bachelor followers seemed to produce them more often in affiliative and resting contexts. This latter indicates that, for bachelor followers, derived calls may serve a particularly important function in bond maintenance. For followers, actively maintaining affiliative relationships with their bachelor leader may be essential, as the leader may both allow them entrance into a reproductive unit following a successful cooperative takeover, and provide them with the opportunity to share in reproductive success (Snyder-Mackler et al., 2012a).

For leaders, on the other hand, derived call production may serve not only to support coalitionary bonding (Dal Pesco et al. 2022; King et al. 2008), but also to facilitate leadership tasks such as initiating travel or managing conflicts (Gustison et al. 2012; Fichtel et al. 2011). For example, reproductive unit leader males have been described to direct calls at their females after conflicts ‘to keep the peace’ (Gustison et al. 2012), which could potentially also explain higher vocal complexity in the agonistic context by the leader males in our study. Unfortunately, in our study we did not annotate whether derived call use in agonistic contexts reflected causes or consequences of aggression, which would be needed to differentiate between these two alternatives. Lastly, primate leaders are often responsible for initiating and coordinating group travel (Fichtel et al. 2011). Indeed, in hamadryas baboons, leader males have the greatest influence in the travel decision-making process (Kummer 1968). In both chacma baboons and geladas, however, decision making is shared between the dominant adult males and females (Dunbar 1983; Sueur 2011). Moreover, baboons notably produce only non-derived calls (i.e. grunts, barks and screams) during both travel (e.g. Byrne 1981; Sueur 2011) and agonism (e.g. Cheney et al. 1995; Gustison et al. 2012; Maciej et al. 2013), whereas geladas produce derived calls in both of these contexts.

It warrants further research why gelada leader males are additionally producing derived calls in travel and agonistic contexts, whereas baboons appear to manage travel activities and agonistic situations without the use of additional call types. Future studies should investigate the functions of the individual derived call types, as well as possible derived call combinations (see below), in these specific contexts.

Call combinations in particular, as well as the occurrence of vocal exchanges and call overlap are exciting areas for future research in geladas. Whether geladas combine derived calls in specific patterns, and thus whether they exhibit combinatorial vocal complexity (Bouchet et al. 2013), remains unknown. Considering that many primates combine vocal types (e.g. Candiotti et al. 2012; Leroux & Townsend 2020) to enhance limited vocal repertoires (Coye et al. 2018; Fitch & Hauser 1995), call combinations in particular may help leader males perform their leader role across social contexts. In addition, early work has described the use of vocal exchanges between reproductive unit leaders and females as additional indicators of bond strength in geladas (Richman 1987), suggesting that such exchanges additionally support bonding. In our study, bachelors engaged in such vocal exchanges as well (pers. obs.). However, whether these occur more frequently between male coalition partners and whether they incorporate derived calls in these exchanges needs further investigation. Intriguingly, we also sometimes observed that bachelors overlap each other’s calls (pers. obs.). This behaviour is also seen in birds during male–male contests, and can signal aggression, advertise quality, or mask rivals (Helfer & Osiejuk 2014; Logue 2021; Naguib & Mennill 2010). In geladas, another bachelor male strategy, besides merely producing more complex sequences to gain dominance status in the bachelor group, could thus be to overlap the complex sequences of rival males. Such exchange and overlap dynamics are thus an interesting additional topic for future research into sexual selection and male quality.

In sum, until now, it has remained debated why geladas evolved a larger vocal repertoire than the closely related baboons (Gustison & Bergman 2017; Hammerschmidt & Fischer 2019; Maciej et al. 2013). This question is particularly relevant for the comparison between geladas, and Guinea and hamadryas baboons, who share similar multi-level social organisations, but who lack the extended vocal repertoire found in geladas (Hammerschmidt & Fischer 2019). Our study suggests that the evolution of derived gelada calls might also lie in the long-term relationships males have to form with each other during their lifetime. Hamadryas and Guinea baboons do not form distinct all-male groups. Instead, bachelor males in these two species only associate with leader males of reproductive units, who also often are relatives (Chowdhury et al. 2015; Dal Pesco et al. 2022). Contrastingly, gelada bachelor groups consist of largely unrelated males (Pappano 2013) that are also potential future rivals. These bachelor males likely form enduring relationships, as they remain in bachelor groups for three to five years (Pappano 2013). Moreover, this bond seems to transfer to reproductive units after a cooperative takeover, as followers often help reproductive unit leaders defend their harem during takeovers, and potentially get to reproduce in return (Snyder-Mackler et al. 2012a; but see Miller et al. 2021). Thus, although Guinea and hamadryas baboons and geladas share a multi-level social system, they differ in the social dynamics among bachelor males. The extended vocal repertoire of geladas may support this interplay between male-male cooperation and competition, which are seen as the drivers of social cognition (Seyfarth & Cheney 2015). This complex interplay may be what sets gelada males apart from *Papio* spp., providing a possible explanation for the evolution of the unique vocal repertoire of geladas.

In conclusion, the results from our pilot study support our hypotheses that derived calls function in both status signalling and bond maintenance, while revealing that leader males use derived calls more broadly than predicted. Taken together, these findings indicate that derived calls produced by gelada males serve functions beyond male-female interactions. This is the first study to have examined the vocal behaviour within gelada bachelor groups. Consequently, many future research opportunities into gelada vocal behaviour are available, as questions remain about the functions of the specific derived call types, as well as indicators of quality related to derived call production, derived call combinations, and vocal exchange dynamics. Moreover, derived calls might play an important role in the formation of bachelor groups. These unexplored research topics into the function of derived gelada calls can help untangle their evolution, as well as provide insight into human language evolution, highlighting that geladas remain an intriguing model species for investigating how vocal complexity functions within socially complex species.

## Statements and declarations

The authors have no relevant financial or non-financial interests to disclose.

Data availability: raw data used for the statistical analyses are available in Supplementary Material 2.

## Acknowledgements

We thank Dr. Tom Roth for statistical advice. We thank Rotterdam Zoo for making this research possible.

## References

Agbangba, C. E., Aide, E. S., Honfo, H., & Kakai, R. G. (2024). On the use of post-hoc tests in environmental and biological sciences: A critical review. Heliyon, 10(3), e25131. 10.1016/j.heliyon.2024.e25131

Altmann, J. (1974). Observational Study of Behavior: Sampling Methods. Behaviour, 49(3–4), 227–266. 10.1163/156853974X00534

Benítez, M. E., le Roux, A., Fischer, J., Beehner, J. C., & Bergman, T. J. (2016). Acoustic and Temporal Variation in Gelada (Theropithecus gelada) Loud Calls Advertise Male Quality. International Journal of Primatology, 37(4–5), 568–585. 10.1007/s10764-016-9922-0

Benítez, M. E., Pappano, D. J., Beehner, J. C., & Bergman, T. J. (2017). Evidence for mutual assessment in a wild primate. Scientific Reports, 7(1), 2952. 10.1038/s41598-017-02903-w

Bergman, T. J. (2013). Speech-like vocalized lip-smacking in geladas. Current Biology, 23(7), R268–R269. 10.1016/j.cub.2013.02.038

Boersma, P., & Weenink, D. (2024). Praat: doing phonetics by computer (6.4.13) [Software]. https://praat.org

Bouchet, H., Blois-Heulin, C., & Lemasson, A. (2013). Social complexity parallels vocal complexity: a comparison of three non-human primate species. Frontiers in Psychology, 4. 10.3389/fpsyg.2013.00390

Brooks, M. E., Kristensen, K., Benthem, K. J., van, Magnusson, A., Berg, C. W., Nielsen, A., Skaug, H. J., Mächler, M., & Bolker, B. M. (2017). glmmTMB Balances Speed and Flexibility Among Packages for Zero-inflated Generalized Linear Mixed Modeling. The R Journal, 9(2), 378. 10.32614/RJ-2017-066

Byrne, R.W. (1981). Distance Vocalisations of Guinea Baboons (Papio papio) in Senegal: An Analysis of Function. Behaviour, 78(3/4), 283–313.

Byrne, R. W. (2016). *Evolving Insight*. Oxford University Press. 10.1093/acprof:oso/9780198757078.001.0001

Candiotti, A., Zuberbühler, K., & Lemasson, A. (2012). Context-related call combinations in female Diana monkeys. Animal Cognition, 15(3), 327–339. 10.1007/s10071-011-0456-8

Cheney, D. L., Seyfarth, R. M., & Silk, J. B. (1995). The role of grunts in reconciling opponents and facilitating interactions among adult female baboons. Animal Behaviour, 50(1), 249–257. 10.1006/anbe.1995.0237

Chowdhury, S., Pines, M., Saunders, J., & Swedell, L. (2015). The adaptive value of secondary males in the polygynous multi level society of hamadryas baboons. American Journal of Physical Anthropology, 158(3), 501–513. 10.1002/ajpa.22804

Coye, C., Ouattara, K., Arlet, M. E., Lemasson, A., & Zuberbühler, K. (2018). Flexible use of simple and combined calls in female Campbell’s monkeys. Animal Behaviour, 141, 171–181. 10.1016/j.anbehav.2018.05.014

Dal Pesco, F., Trede, F., Zinner, D., & Fischer, J. (2022). Male–male social bonding, coalitionary support and reproductive success in wild Guinea baboons. Proceedings of the Royal Society B: Biological Sciences, 289(1975). 10.1098/rspb.2022.0347

Dunbar, R. I. M. (1980). Demographic and Life History Variables of a Population of Gelada Baboons (Theropithecus gelada). Journal of Animal Ecology, 49(2), 485–506. 10.2307/4259

Dunbar, R. I. M. (1983). Structure of Gelada Baboon Reproductive Units. IV. Integration at Group Level. Zeitschrift Für Tierpsychologie, 63(4), 265–282. 10.1111/j.1439-0310.1983.tb00743.x

Dunbar, R.I.M. (1984). *Reproductive Decisions: An Economic Analysis of Gelada Baboon Social Strategies*. Princeton University Press.

Farine, D. R., & Sanchez-Tojar, A. (2021). aniDom: Inferring Dominance Hierarchies and Estimating Uncertainty (0.1.5).

Fashing, P.J., Nguyen, N., Venkataraman, V.V., & Kerby, J.T. (2014). Gelada feeding ecology in an intact ecosystem at Guassa, Ethiopia: Variability over time and implications for theropith and hominin dietary evolution. American Journal of Physical Anthropology, 155(1), 1–16. 10.1002/ajpa.22559

Fichtel, C., Pyritz, L., & Kappeler, P. M. (2011). Coordination of Group Movements in Non-human Primates. In Coordination in Human and Primate Groups (pp. 37–56). Springer Berlin Heidelberg. 10.1007/978-3-642-15355-6_3

Fischer, J., Kitchen, D. M., Seyfarth, R. M., & Cheney, D. L. (2004). Baboon loud calls advertise male quality: acoustic features and their relation to rank, age, and exhaustion. Behavioral Ecology And Sociobiology, 56(2), 140–148. 10.1007/s00265-003-0739-4

Fitch, W. T. (1997). Vocal tract length and formant frequency dispersion correlate with body size in rhesus macaques. The Journal Of The Acoustical Society Of America, 102(2), 1213–1222. 10.1121/1.421048

Fitch, W. T., & Hauser, M. D. (1995). Vocal production in nonhuman primates: Acoustics, physiology, and functional constraints on “honest” advertisement. American Journal of Primatology, 37(3), 191–219. 10.1002/ajp.1350370303

Fitch, W. T., & Hauser, M. D. (2003). Unpacking “Honesty”: Vertebrate Vocal Production and the Evolution of Acoustic Signals. In Acoustic Communication (pp. 65–137). Springer-Verlag. 10.1007/0-387-22762-8_3

Freeberg, T. M. (2006). Social Complexity Can Drive Vocal Complexity. Psychological Science, 17(7), 557–561. 10.1111/j.1467-9280.2006.01743.x

Freeberg, T. M., Dunbar, R. I. M., & Ord, T. J. (2012). Social complexity as a proximate and ultimate factor in communicative complexity. Philosophical Transactions of the Royal Society B: Biological Sciences, 367(1597), 1785–1801. 10.1098/rstb.2011.0213

Galbany, J., Tung, J., Altmann, J., & Alberts, S. C. (2015). Canine Length in Wild Male Baboons: Maturation, Aging and Social Dominance Rank. PLOS ONE, 10(5), e0126415. 10.1371/journal.pone.0126415

Gippoliti, S., Mekonnen, A., Burke, R., Nguyen, N., & Fashing, P.J. (2025). *Theropithecus gelada* (amended version of 2022 assessment). The IUCN Red List of Threatened Species 2025: e.T21744A274974485. 10.2305/IUCN.UK.2025-1.RLTS.T21744A274974485.en.

Gustison, M. L., & Bergman, T. J. (2016). Vocal complexity influences female responses to gelada male calls. Scientific Reports, 6(1), 19680. 10.1038/srep19680

Gustison, M. L., & Bergman, T. J. (2017). Divergent acoustic properties of gelada and baboon vocalizations and their implications for the evolution of human speech. Journal of Language Evolution, 2(1), 20–36. 10.1093/jole/lzx015

Gustison, M. L., le Roux, A., & Bergman, T. J. (2012). Derived vocalizations of geladas (Theropithecus gelada) and the evolution of vocal complexity in primates. Philosophical Transactions of the Royal Society B: Biological Sciences, 367(1597), 1847–1859. 10.1098/rstb.2011.0218

Gustison, M. L., Semple, S., Ferrer-I-Cancho, R., & Bergman, T. J. (2016). Gelada vocal sequences follow Menzerath’s linguistic law. Proceedings Of The National Academy Of Sciences, 113(19), E2750–E2758. 10.1073/pnas.1522072113

Gustison, M. L., Tinsley Johnson, E., Beehner, J. C., & Bergman, T. J. (2019). The social functions of complex vocal sequences in wild geladas. Behavioral Ecology and Sociobiology, 73(1), 14. 10.1007/s00265-018-2612-5

Hammerschmidt, K., & Fischer, J. (2019). Baboon vocal repertoires and the evolution of primate vocal diversity. Journal of Human Evolution, 126, 1–13. 10.1016/j.jhevol.2018.10.010

Hartig, F. (2022). DHARMa: Residual Diagnostics for Hierarchical (Multi-Level / Mixed) Regression Models (0.4.6).

Helfer, B., & Osiejuk, T. S. (2015). It Takes All Kinds in Acoustic Communication: A New Perspective on the Song Overlapping Phenomenon. Ethology, 121(4), 315–326. 10.1111/eth.12356

Hewitt, G., MacLarnon, A., & Jones, K. E. (2002). The Functions of Laryngeal Air Sacs in Primates: A New Hypothesis. Folia Primatologica, 73(2–3), 70–94. 10.1159/000064786

Hohn, T. I., Lin, B., Miller, C. M., Foxfoot, I. R., Venkataraman, V. V., Ruckstuhl, K. E., Nguyen, N., & Fashing, P. J. (2024). Post-Conflict Behaviors of Wild Gelada Monkeys (Theropithecus gelada) at Guassa, Ethiopia. International Journal Of Primatology, 45(5), 1083–1106. 10.1007/s10764-024-00438-2

Hotchkin, C., & Parks, S. (2013). The Lombard effect and other noise induced vocal modifications: insight from mammalian communication systems. Biological Reviews, 88(4), 809–824. 10.1111/brv.12026

Kappeler, P. M. (2019). A framework for studying social complexity. Behavioral Ecology and Sociobiology, 73(1), 13. 10.1007/s00265-018-2601-8

Kavanagh, E., Street, S. E., Angwela, F. O., Bergman, T. J., Blaszczyk, M. B., Bolt, L. M., Briseño-Jaramillo, M., Brown, M., Chen-Kraus, C., Clay, Z., Coye, C., Thompson, M. E., Estrada, A., Fichtel, C., Fruth, B., Gamba, M., Giacoma, C., Graham, K. E., Green, S., . . . Slocombe, K. (2021). Dominance style is a key predictor of vocal use and evolution across nonhuman primates. Royal Society Open Science, 8(7), 210873. 10.1098/rsos.210873

Kawai, M., Ohsawa, H., Mori, U., & Dunbar, R. (1983). Social organization of gelada baboons: Social units and definitions. Primates, 24(1), 13–24. 10.1007/BF02381450

King, A. J., Douglas, C. M. S., Huchard, E., Isaac, N. J. B., & Cowlishaw, G. (2008). Dominance and Affiliation Mediate Despotism in a Social Primate. Current Biology, 18(23), 1833–1838. 10.1016/j.cub.2008.10.048

Knörnschild, M., Fernandez, A. A., & Nagy, M. (2020). Vocal information and the navigation of social decisions in bats: Is social complexity linked to vocal complexity? Functional Ecology, 34(2), 322–331. 10.1111/1365-2435.13407

Krams, I., Krama, T., Freeberg, T. M., Kullberg, C., & Lucas, J. R. (2012). Linking social complexity and vocal complexity: a parid perspective. Philosophical Transactions of the Royal Society B: Biological Sciences, 367(1597), 1879– 1891. 10.1098/rstb.2011.0222

Kummer, H. (1968). *Social organization of hamadryas baboons: A field study*. The University of Chicago Press.

Lenth, R. v. (2024). emmeans: Estimated Marginal Means, aka Least-Squares Means (1.10.1).

Leone, A., & Palagi, E. (2010). Reconciling conflicts in a one-male society: the case of geladas (Theropithecus gelada). Primates, 51(3), 203–212. 10.1007/s10329-010-0188-4

Leroux, M., & Townsend, S. W. (2020). Call combinations in great apes and the evolution of syntax. Animal Behavior and Cognition, 7(2), 131–139. 10.26451/abc.07.02.07.2020

Lincoln Park Zoo. (2022). ZooMonitor (4.1).

Logue, D. M. (2021). Countersinging in birds (pp. 1–61). 10.1016/bs.asb.2021.03.001

Lüdecke, D. (2018). ggeffects: Tidy Data Frames of Marginal Effects from Regression Models. Journal of Open Source Software, 3(26), 772. 10.21105/joss.00772

Lüdecke, D., Ben-Shachar, M., Patil, I., Waggoner, P., & Makowski, D. (2021). performance: An R Package for Assessment, Comparison and Testing of Statistical Models. Journal of Open Source Software, 6(60), 3139. 10.21105/joss.03139

Maciej, P., Ndao, I., Hammerschmidt, K., & Fischer, J. (2013). Vocal communication in a complex multi-level society: constrained acoustic structure and flexible call usage in Guinea baboons. Frontiers in Zoology, 10(1), 58. 10.1186/1742-9994-10-58

Manser, M. B., Jansen, D. A. W. A. M., Graw, B., Hollén, L. I., Bousquet, C. A. H., Furrer, R. D., & le Roux, A. (2014). Vocal Complexity in Meerkats and Other Mongoose Species. In M. Naguib, L. Barrett, H. J. Brockmann, S. Healy, J. C. Mitani, T. J. Roper, & L. W. Simmons (Eds.), Advances in the Study of Behavior (Vol. 46, pp. 281–310). Academic Press. 10.1016/B978-0-12-800286-5.00006-7

McComb, K., & Semple, S. (2005). Coevolution of vocal communication and sociality in primates. Biology Letters, 1(4), 381–385. 10.1098/rsbl.2005.0366

Miller, C. M., Snyder-Mackler, N., Nguyen, N., Fashing, P. J., Tung, J., Wroblewski, E. E., Gustison, M. L., & Wilson, M. L. (2021). Extragroup paternity in gelada monkeys, Theropithecus gelada, at Guassa, Ethiopia and a comparison with other primates. Animal Behaviour, 177, 277–301. 10.1016/j.anbehav.2021.05.008

Naguib, M., & Mennill, D. J. (2010). The signal value of birdsong: empirical evidence suggests song overlapping is a signal. Animal Behaviour, 80(3), e11–e15. 10.1016/j.anbehav.2010.06.001

Neumann, C., Duboscq, J., Dubuc, C., Ginting, A., Irwan, A. M., Agil, M., Widdig, A., & Engelhardt, A. (2011). Assessing dominance hierarchies: validation and advantages of progressive evaluation with Elo-rating. Animal Behaviour, 82(4), 911–921. 10.1016/j.anbehav.2011.07.016

Palagi, E., Leone, A., Demuru, E., & Ferrari, P. F. (2018). High-Ranking Geladas Protect and Comfort Others After Conflicts. Scientific Reports, 8(1), 15291. 10.1038/s41598-018-33548-y

Pappano, D. J. (2013). *The Reproductive Trajectories of Bachelor Geladas*. https://deepblue.lib.umich.edu/handle/2027.42/102358

Plavcan, J. M. (2012). Sexual Size Dimorphism, Canine Dimorphism, and Male-Male Competition in Primates. Human Nature, 23(1), 45–67. 10.1007/s12110-012-9130-3

Puts, D. A., Hill, A. K., Bailey, D. H., Walker, R. S., Rendall, D., Wheatley, J. R., Welling, L. L. M., Dawood, K., Cárdenas, R., Burriss, R. P., Jablonski, N. G., Shriver, M. D., Weiss, D., Lameira, A. R., Apicella, C. L., Owren, M. J., Barelli, C., Glenn, M. E., & Ramos-Fernandez, G. (2016). Sexual selection on male vocal fundamental frequency in humans and other anthropoids. Proceedings of the Royal Society B: Biological Sciences, 283(1829), 20152830. 10.1098/rspb.2015.2830

R Core Team. (2024). R: A Language and Environment for Statistical Computing (4.4.0).

Richman, B. (1987). Rhythm and melody in gelada vocal exchanges. Primates, 28(2), 199–223. 10.1007/BF02382570

Seyfarth, R. M., & Cheney, D. L. (2015). Social cognition. Animal Behaviour, 103, 191–202. 10.1016/j.anbehav.2015.01.030

Snyder-Mackler, N., Alberts, S. C., & Bergman, T. J. (2012a). Concessions of an alpha male? Cooperative defence and shared reproduction in multi-male primate groups. Proceedings of the Royal Society B: Biological Sciences, 279(1743), 3788–3795. 10.1098/rspb.2012.0842

Snyder-Mackler, N., Beehner, J. C., & Bergman, T. J. (2012b). Defining Higher Levels in the Multilevel Societies of Geladas (Theropithecus gelada). International Journal of Primatology, 33(5), 1054–1068. 10.1007/s10764-012-9584-5

Soma, M., & Garamszegi, L. Z. (2011). Rethinking birdsong evolution: meta-analysis of the relationship between song complexity and reproductive success. Behavioral Ecology, 22(2), 363–371. 10.1093/beheco/arq219

Sueur, C. (2011). Group decision-making in chacma baboons: leadership, order and communication during movement. BMC Ecology, 11(1), 26. 10.1186/1472-6785-11-26

Swedell, L., & Schreier, A. (2009). Male aggression towards females in Hamadryas Baboons: conditioning, coercion and control. In M. N. Muller & R. W. Wrangham (Eds.), Sexual Coercion in Primates and Humans: An Evolutionary Perspective on Male Aggression Against Females (pp. 244–270). Harvard University Press.

The Audacity Team. (2024). *Audacity* (3.5.1).

Tibbetts, E. A., Pardo-Sanchez, J., & Weise, C. (2022). The establishment and maintenance of dominance hierarchies. Philosophical Transactions Of The Royal Society B Biological Sciences, 377(1845), 20200450. 10.1098/rstb.2020.0450

Tinsley Johnson, E., Snyder-Mackler, N., Beehner, J. C., & Bergman, T. J. (2014). Kinship and Dominance Rank Influence the Strength of Social Bonds in Female Geladas (Theropithecus gelada). International Journal of Primatology, 35(1), 288–304. 10.1007/s10764-013-9733-5

Titze, I. R., & Riede, T. (2010). A Cervid Vocal Fold Model Suggests Greater Glottal Efficiency in Calling at High Frequencies. PLoS Computational Biology, 6(8), e1000897. 10.1371/journal.pcbi.1000897

Wickham, H. (2016). ggplot2: Elegant Graphics for Data Analysis. Springer-Verlag New York.

Wright, E., Galbany, J., McFarlin, S. C., Ndayishimiye, E., Stoinski, T. S., & Robbins, M. M. (2019). Male body size, dominance rank and strategic use of aggression in a group-living mammal. Animal Behaviour, 151, 87–102. 10.1016/j.anbehav.2019.03.011

Zinner, D., Atickem, A., Beehner, J. C., Bekele, A., Bergman, T. J., Burke, R., Dolotovskaya, S., Fashing, P. J., Gippoliti, S., Knauf, S., Knauf, Y., Mekonnen, A., Moges, A., Nguyen, N., Stenseth, N. C., & Roos, C. (2018). Phylogeography, mitochondrial DNA diversity, and demographic history of geladas (Theropithecus gelada). PLoS ONE, 13(8), e0202303. 10.1371/journal.pone.0202303

