## Supplementary Material 1 for "Beyond the harem: vocal complexity in a captive gelada (*Theropithecus gelada*) bachelor group"

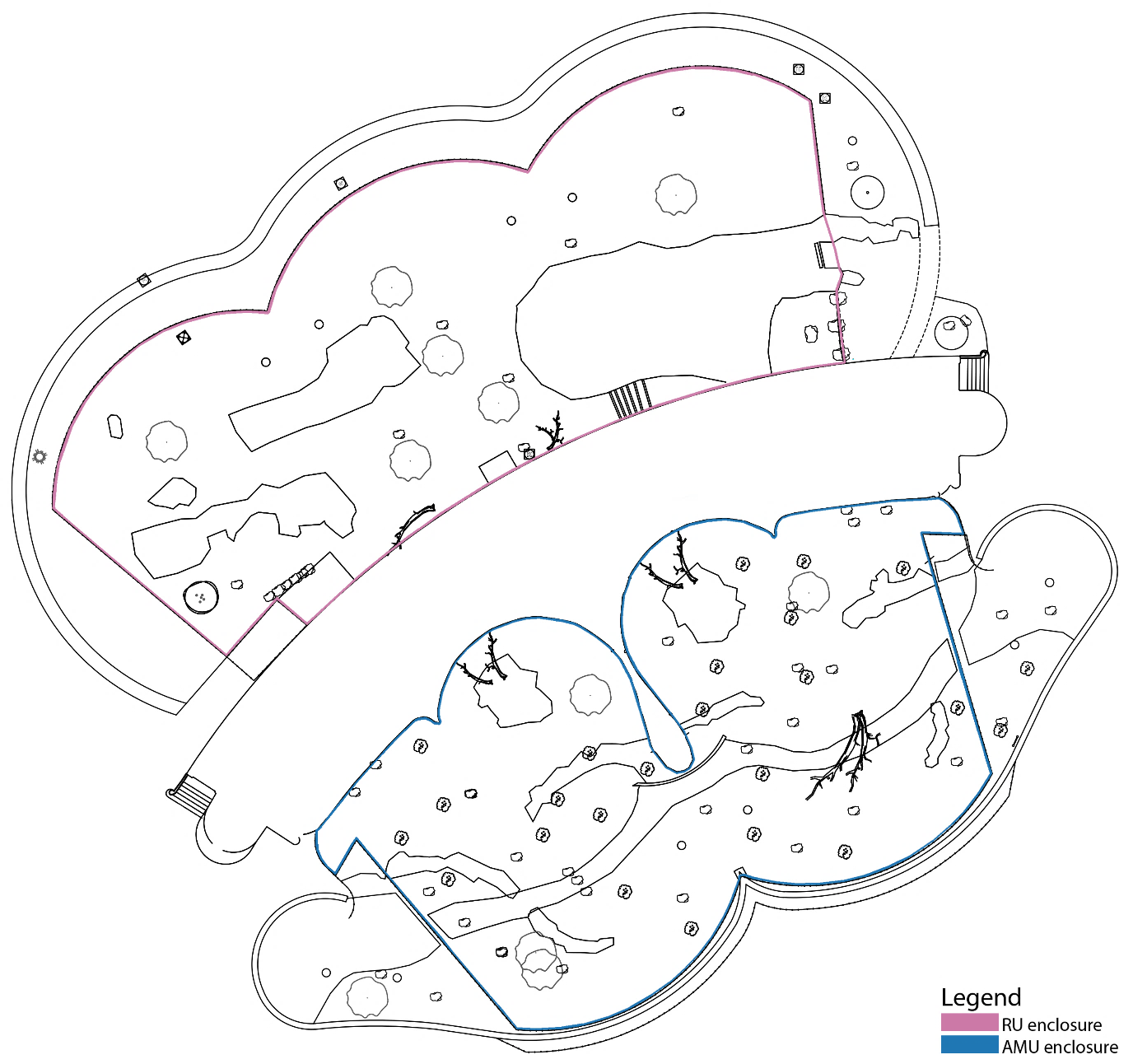

**Figure S1.** Map of the outside enclosures of the reproductive unit (pink outline) and all-male unit (blue outline). Between the enclosure is the building housing the inside enclosures, caretaker area and visitor area

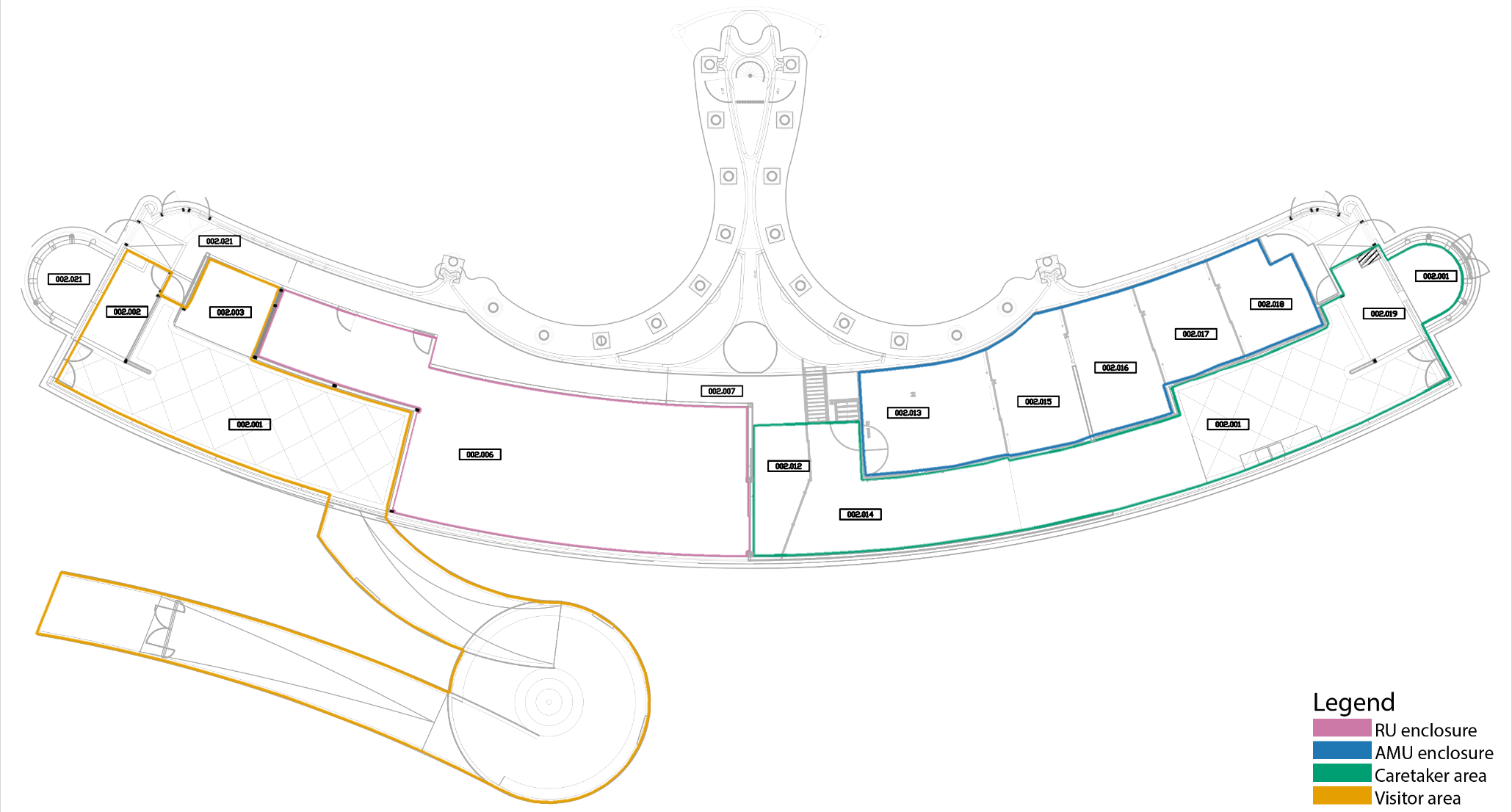

**Figure S2.** Map of the building containing the inside enclosure of the reproductive unit (pink outline) and all-male unit (blue outline), and caretaker area (green outline) and visitor area (yellow outline)

**Table S1.** Ethogram of the social behaviours of geladas used to determine the contextual categories of the vocalisations

| Category | Behaviour | Description |
| --- | --- | --- |
| Affiliation | Approach | An individual calmly moves towards another individual in a non-aggressive manner seemingly with the goal of social interaction (Taberer et al. 2022). |
|  | Grooming | An individual is moving and picking its hands through the fur or picking the skin of another individual while moving its hands to its own mouth occasionally (Alvarez & Cónsul 1978). |
|  | Lip-smacking | An individual rapidly opens and closes its lips repetitively, allowing the tongue to protrude (Alvarez & Cónsul 1978; Maestripieri & Wallen 1997). Can occur together with a wobble (Bergman 2013; Gustison et al. 2012). |
|  | Contact | An individual is sitting in physical contact with another individual. |
|  | Proximity | An individual is sitting within two arm’s length next to another individual. |
|  | Withdraw | An individual moves out of contact from another individual (Carter 2016). |
| Agonism | Threat | An individual holds its gaze towards another individual including the raising of eyebrows, retraction of the scalp and moving back of the ears (Taberer et al. 2022). |
|  | Chase | An individual is hastily running after another individual, includes lunging forward to another individual (Taberer et al. 2022). |
|  | Aggression | An individual is, for example, biting, hitting, pushing or grabbing another individual (Taberer et al. 2022). |
|  | Eyebrow raise | An individual moves its eyebrows up and down such that the light skin above the eyes becomes tense and more visible (Taberer et al. 2022). |
|  | Displacement | An individual approaches another individual, forcing it to move, and/or usurping its foraging spot or grooming interaction (Le Roux et al. 2011). |
|  | Give ground | An individual increases the distance from an approaching individual by walking, running or jumping away (Alvarez & Cónsul 1978). |
|  | Bare teeth | The mouth of an individual is closed, or slightly open, but the lips are retracted, exposing the teeth (Maestripieri & Wallen 1997). The mouth can also move rapidly open and closed (Alvarez & Cónsul 1978). |
|  | Agonism response | An individual is, for example, looking at and moving around an occurring conflict and responding to this conflict by vocalizing (Gustison et al. 2012). |
| Foraging | Foraging | The focal individual is plucking grass or hay, digging in the soil, or eating vegetables (Filipčík et al. 2014). |
| Rest | Sitting | An individual is sitting, but is vigilant, its eyes are open, looking around in a relaxed manner. |
|  | Resting | The focal individual is sitting or lying down with no additional activities (Filipčík et al. 2014). Its head is down and/or its eyes are closed. |
| Travelling | Moving | The focal individual is moving by walking, running or climbing. |
|  | Follow | An individual is walking, running or climbing in the same direction as another individual (Taberer et al. 2022). |
|  | Travel response | An individual vocally responds to its group members moving or following each other. |

References Table S1

Alvarez, F., & Cónsul, C. (1978). The structure of social behaviour in Theropithecus gelada. *Primates*, *19*(1), 45–59.	<https://doi.org/10.1007/BF02373226>

Bergman, T. J. (2013). Speech-like vocalized lip-smacking in geladas. *Current Biology*, *23*(7), R268–R269.	<https://doi.org/10.1016/j.cub.2013.02.038>

Carter, A. R. (2016). *Variation in patterns of allocare in captive Hamadryas baboons (Papio hamadryas): The potential	effects of environment and kinship in the development of novel behavior*.

Filipčík, R., Máchal, L., Baholet, D., Chládek, G., Hošek, M., & Paldusová, M. (2014). The Daily Pattern of Main Activities	in the Gelada Baboon (Theropithecus gelada). *Acta Universitatis Agriculturae et Silviculturae Mendelianae	Brunensis*, *62*(5), 891–896. <https://doi.org/10.11118/actaun201462050891>

Gustison, M. L., le Roux, A., & Bergman, T. J. (2012). Derived vocalizations of geladas ( Theropithecus gelada ) and the	evolution of vocal complexity in primates. *Philosophical Transactions of the Royal Society B: Biological Sciences*,	*367*(1597), 1847–1859. <https://doi.org/10.1098/rstb.2011.0218>

le Roux, A., Beehner, J. C., & Bergman, T. J. (2011). Female philopatry and dominance patterns in wild geladas. *American	Journal of Primatology*, *73*(5), 422–430. <https://doi.org/10.1002/ajp.20916>

Maestripieri, D., & Wallen, K. (1997). Affiliative and submissive communication in rhesus macaques. *Primates*, *38*(2), 127–	138. <https://doi.org/10.1007/BF02382003>

Taberer, T. R., Mead, J., Hartley, M., & Harvey, N. D. (2023). Impact of female contraception for population management	on behavior and social interactions in a captive troop of Guinea baboons ( *Papio papio* ). *Zoo Biology*, *42*(2), 254–	267. <https://doi.org/10.1002/zoo.21728>
